# T-cell activation is modulated by the 3D mechanical microenvironment

**DOI:** 10.1101/580886

**Authors:** Fatemeh S. Majedi, Mohammad Mahdi Hasani-Sadrabadi, Timothy J. Thauland, Song Li, Louis-S. Bouchard, Manish J. Butte

## Abstract

T cells recognize mechanical forces through a variety of cellular pathways, including mechanical triggering of the T-cell receptor (TCR) and mechanical triggering of the integrin LFA-1. We show here that T cells can recognize forces arising from the rigidity of the microenvironment. We fabricated 3D hydrogels with mechanical stiffness tuned to 4 kPa and 40 kPa and specially engineered be microporous independent of stiffness. We cultured T cells and antigen presenting cells within the matrices and studied activation by flow cytometry and live imaging. We found there was an augmentation of T-cell activation in the context of mechanically stiffer 3D material as compared to the softer material. In contrast, proliferation, activation markers, and migration were all diminished in T cells cultured in the softer material. These results show that T cells can sense their mechanical environment and amplify responses in the context of mechanical stiffness.

## Introduction

When T cells recognize their cognate antigen on the surface of antigen presenting cells (APCs), they form a complex three-dimensional structure known as the immune synapse (IS) (Dustin, 2011). The IS facilitates communication between T cells and APCs via receptor-ligand interactions, is critical for T-cell activation, and is a platform for the delivery of effector molecules (e.g. the contents of cytolytic granules). At the molecular level, receptor-ligand interactions at the IS trigger kinase-mediated signaling cascades and the release of calcium from intracellular stores, and these TCR-proximal signals lead to the initiation of transcriptional programs controlling T-cell proliferation, differentiation, and effector function.

The actin cytoskeleton is critical for T-cell biology, and pharmacological disruption of F-actin severely cripples T-cell motility, IS formation, and T-cell activation (Burkhardt et al., 2008). While cytoskeletal rearrangement is critical for IS formation, the state of the cytoskeleton prior to T cell-APC conjugation is also important. We have previously shown that naive T cells are mechanically stiffer than activated effector T cells (Thauland et al., 2017). This difference in stiffness allows the more pliable effector T cells to form larger IS with APCs, increasing the number of receptor-ligand interactions and enhancing activation. Thus, the T-cell cytoskeleton acts as a layer of regulation, allowing effector cells to respond to APCs with exquisite sensitivity. The stiffness of the antigen-presenting surface that T cells interact with also plays a role in activation. Experiments examining the interaction of T cells with stimulatory 2D surfaces of varying stiffness have demonstrated that relatively stiffer surfaces provide a stronger stimulus (Judokusumo et al., 2012; O’Connor et al., 2012; Tabdanov et al., 2015; Husson et al., 2011; Hui et al., 2015; Saitakis et al., 2017). The cytotoxicity of T cells is also affected by substrate stiffness (Basu et al., 2016).

While the mechanical properties of the T cell and the antigen-presenting surface have been investigated, the effect of the mechanical properties of the 3D microenvironment on actin-dependent T-cell functions such as motility, IS formation, and activation has not been examined. The properties of 3D hydrogels can be manipulated by altering the density of polymers and crosslinkers. As cells migrate through hydrogels, they interact with functional groups, that act as ligands for cell surface receptors, presented by the polymer. Thus, increasing polymer density results in a concomitant increase in ligand density and a decrease in mesh size, which alters cell-polymer interactions. On the other hand, keeping polymer content constant and varying the crosslinker density also alters the mesh size of the gel, which directly impacts molecular diffusion (Vining and Mooney, 2017). While stiffness and ligand density can be modulated independently of polymer content with some synthetic polymers, the direct relationship between crosslinker density and mesh size makes it difficult to relate cell behavior to pure stiffness. Fortunately, in the case of the biodegradable polymer alginate gelation is induced by calcium ions. Due to the formation of G-blocks that provide pockets for calcium entrapment, the stiffness of the gel can be altered by varying the concentration of calcium ions without altering the ligand density or pore size.

Here we sought to mimic the range of biologically relevant stiffnesses that T cells experience while patrolling secondary lymphoid organs and peripheral tissues. We developed 3D, porous alginate-based scaffolds that had similar rigidities to lymph nodes. Due to the unique characteristics that the alginate biopolymer possesses, we were able to modulate the rigidity of our scaffolds without affecting their porosity or ligand density. We used these scaffolds to study motility, proliferation, morphology, and immune synapse formation by CD4+ T cells. We found that mechanically stiffer scaffolds enhanced T-cell activation in 3D cultures, akin to what was observed for 2D substrates. Our morphological studies of T cells in hard and soft scaffolds revealed increased cell spreading in hard scaffolds. This effect likely contributes to the enhanced immune synapse size, activation, and proliferation we observed for cells cultured in stiff scaffolds.

## Results and Discussion

To study how the mechanical stiffness of a 3D environment affects T-cell activation, proliferation, and IS formation, we developed macroporous, alginate-based, 3D hydrogels. In order to improve cell adhesion within these scaffolds, the alginate polymer was modified with RGD peptides. We screened polymers with a wide range of mechanical properties by titrating polymer and calcium concentration. Measurements of polymer stiffness demonstrated that we could modulate stiffness over two orders of magnitude by varying alginate density from 0.5-2.5% and calcium concentration from 10-60 mM (**Fig. 1A**). Gels with an order of magnitude difference in stiffness (4 and 40 kPa) in the biologically relevant range were chosen for further study. To verify that changing the stiffness did not result in a change in pore size, we took images of the microporous structure of our 3D gels by scanning electron microscopy (SEM) (**Fig. 1B**). We found that both the 40 and 4 kPa gels had a pore size of approximately 130 microns (**Fig. 1C**).

**Figure 1.**
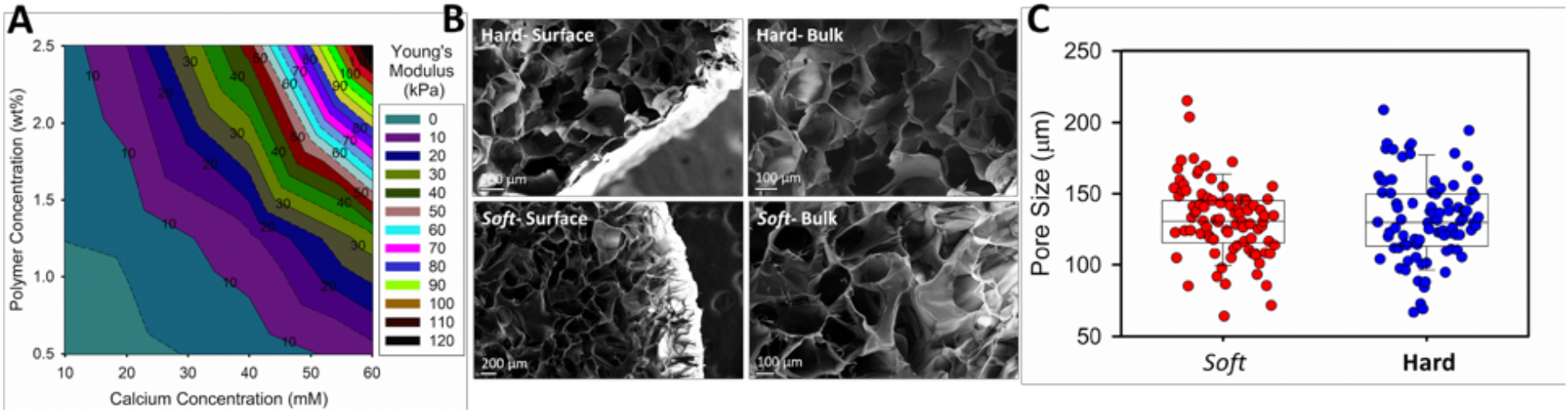
Modulation of alginate gel stiffness without affecting porosity. **(A)** Different formulations of gels were screened by changing the concentration of the alginate polymer or calcium crosslinker. Stiffness values (Young’s modulus) were measured on an Instron mechanical tester. **(B)** SEM images of two gels with an order of magnitude difference in mechanical stiffness. Hard gels were 40 kPa and soft gels were 4 kPa. **(C)** The size of at least 80 pores in each scaffold were measured from SEM images. Average pore size was 131.9 +/− 27.63 μm for hard gels and 132.5 +/− 28 μm for soft gels and was not significantly different.

Given the importance of motility for T-cell function, we attempted to track T-cell migration in hard and soft gels. Fluorescently-labeled cells were seeded into gels and imaged over time. We tracked T cells crawling within the 3D hydrogels (**Fig. 2A**). These data revealed that the average velocity of T cells crawling within hard scaffolds (5.2 μm/min) was significantly higher than within soft scaffolds (4.0 μm/min) (**Fig. 2B**). The higher velocity of cells within stiffer hydrogels most likely leads to a higher probability of interaction between T cells and APCs. Thus, these findings are consistent with the fact that T cells were more activated and proliferative in the stiffer gels.

**Figure 2.**
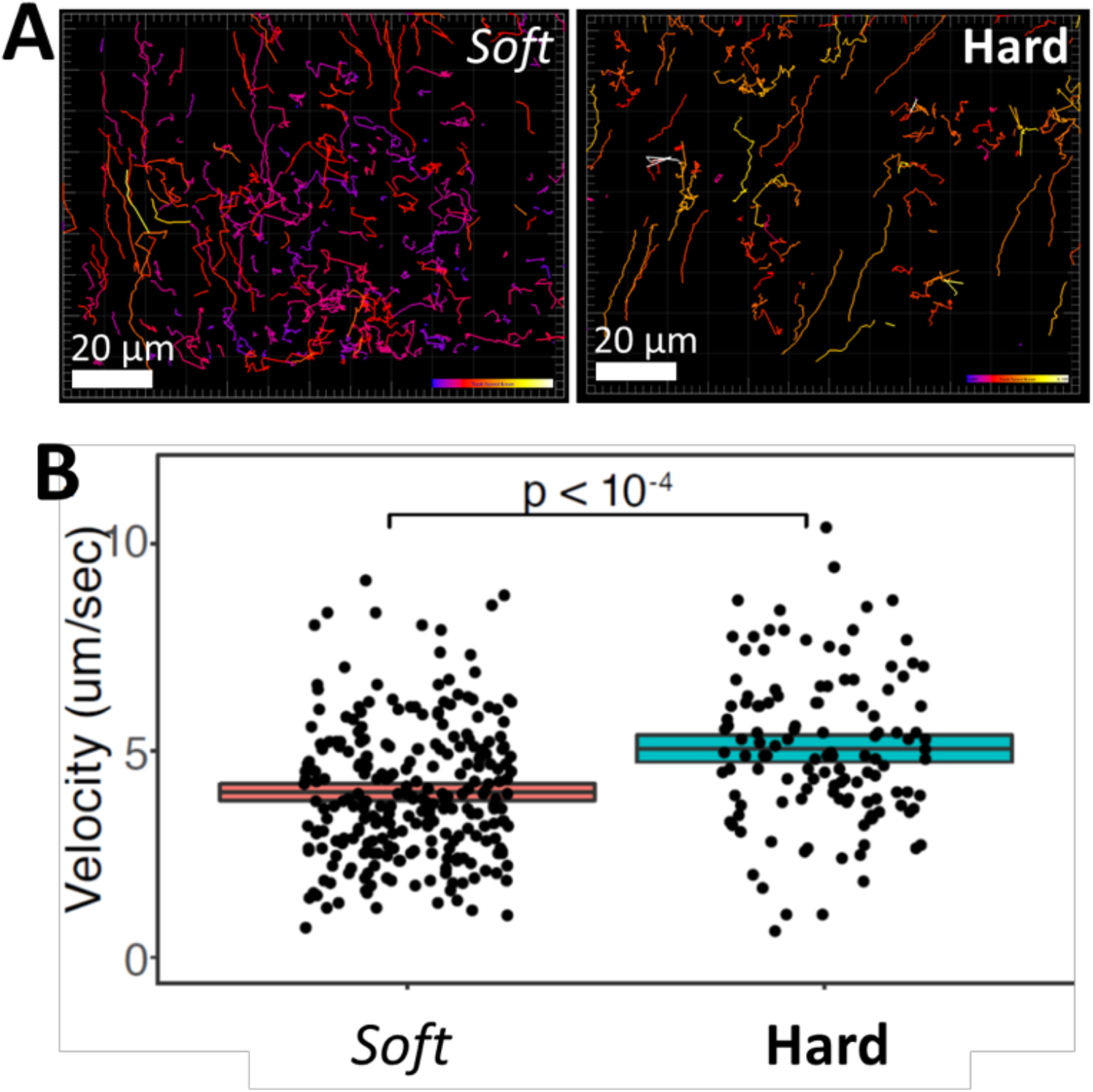
T-cell motility is enhanced in stiff 3D hydrogels. **(A)** T cells crawling through macroporous alginate hydrogels were imaged for 10 min with confocal microscopy. T-cell migration in soft 4 kPa gels (left) and stiff 40 kPa gels (right) were tracked with Imaris. Tracks are color-coded for instantaneous velocity. **(B)** The mean velocity of cells crawling through soft and hard hydrogels is plotted.

To examined the effect of 3D hydrogel stiffness on CD4+ T-cell activation, we cultured T cells and monitored their proliferation. We seeded CFSE-labeled CD4+ T cells purified from OT-II TCR transgenic mice plus peptide-loaded APCs into hard (40 kPa) and soft (4 kPa) 3D alginate scaffolds. To contrast 3D culture with conventional approaches, we prepared 2D alginate-RGD gels with same mechanical stiffnesses. T cells and peptide-loaded APCs were placed onto these 2D substrates. Sequential generations of daughter cells result in roughly two-fold dilution of the CFSE when analyzed by flow cytometry. We also monitored the expression of the activation marker CD25 on the T cells. The stiffer substrate induced more proliferation in both 2D and 3D conditions (**Fig. 3A and B**). Interestingly, the effect of stiffness on proliferation was greater in 3D scaffolds than on 2D (**Fig. 3C and D**). There were slightly more undivided cells in the 3D conditions. We hypothesize that the increased degrees of freedom experienced by cells in a 3D matrix makes successful T cell-APC conjugation less frequent, and thus some T cells remained unactivated, as compared with crawling on a 2D surface. However, the proliferation index – a measure of the proliferative response of the cells that divided at least once – was higher in the stiff 3D environment, suggesting that there was enhanced activation when T cells encounter an APC in 3D (**Fig. 3D**). In agreement with the proliferation data, upregulation of CD25 was more robust on stiffer substrates (**Fig. 3E**).

**Figure 3.**
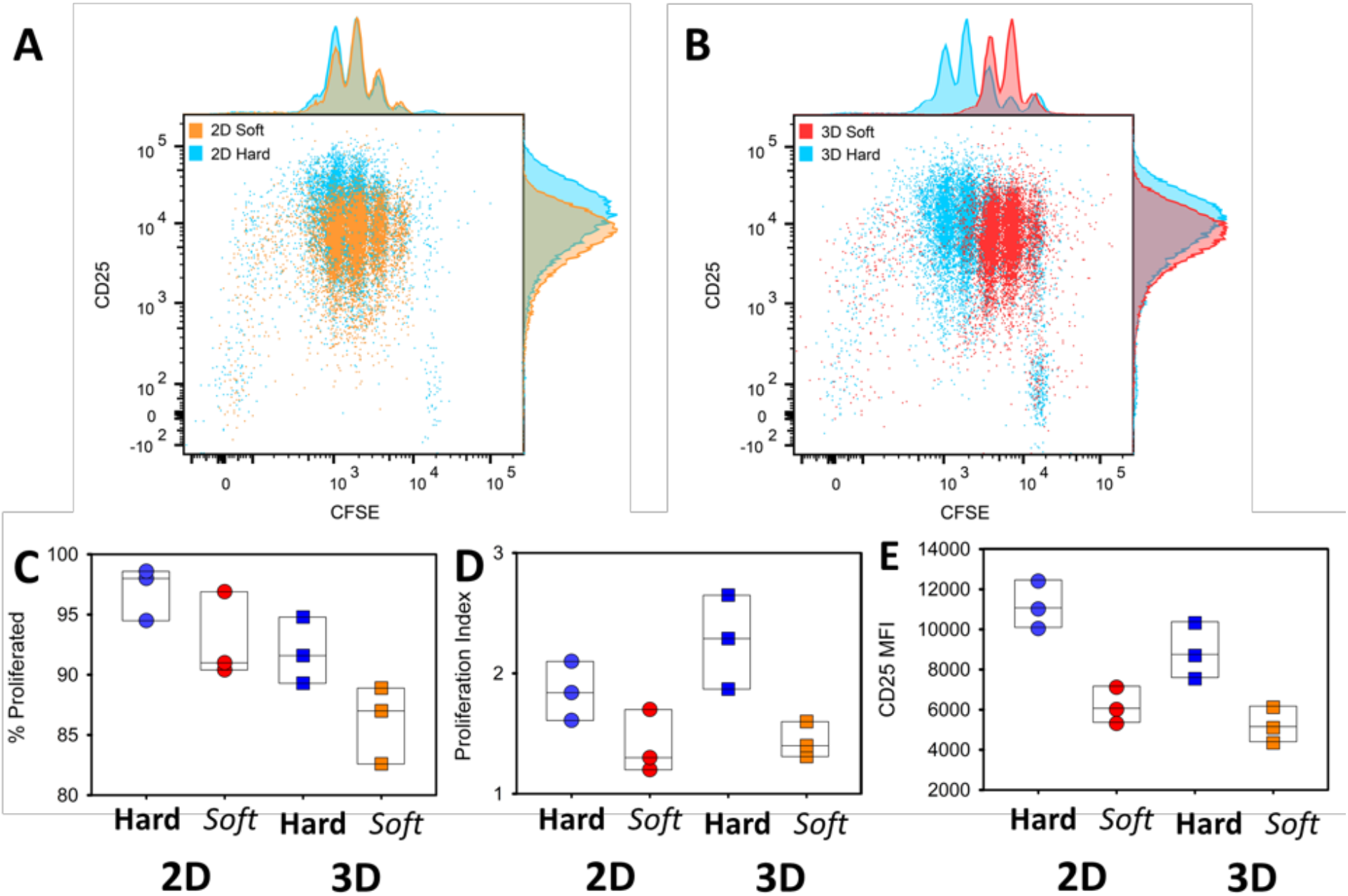
T-cell activation is modulated by the stiffness of 3D hydrogels. (**A** and **B**) FACS analysis of CD25 expression and cell division (CFSE dilution) of CD4+ T cells cocultured with APCs on 2D **(A)** or 3D **(B)** hydrogels with different stiffnesses. Cells were assayed at three days post-stimulation. **(C-E)** Percentage of cells that divided at least once **(C)**, proliferation index of divided cells and mean fluorescence intensity (MFI) of CD25 **(E)** are plotted for cells activated on 2D or 3D and stiff or soft hydrogels. Each dot represents one experiment.

Next, we studied the IS formed by T cells in our 3D alginate hydrogels and compared them to synapses formed on 2D hydrogels. We have previously shown that the size of the IS is correlated with T-cell activation (Thauland et al., 2017). We again seeded CD4+ T cells purified from spleens of OT-II TCR transgenic mice plus ovalbumin-loaded APCs into hard (40 kPa) and soft (4 kPa) 3D alginate scaffolds. To compare synapses formed in 3D with 2D, we prepared 2D alginate-RGD gels of the same mechanical stiffnesses. T cells and peptide-loaded APCs were placed onto these 2D substrates. As a control, we generated T cell-APC synapses by co-culturing them in an Eppendorf tube for 15 min (see Methods). As a proxy for IS size, we measured the volume of the adhesion molecule LFA-1 that accumulated at the T cell-APC interface. LFA-1 plays a crucial role in IS formation (Grakoui et al., 1999), and is not expressed on the B cells that we used as APCs in these experiments, making it an ideal protein to define the extent of the T-cell IS. We also stained the T cell-APC conjugates for the cytoskeletal protein F-actin. We found that immune synapses of T cells formed in stiff matrices (**Fig. 4A**) were significantly larger than those formed in soft matrices (**Fig. 4B**). This consequence was true for T cells activated in 3D matrices or upon 2D substrates (**Fig. 4C**). The IS formed on stiff hydrogels were larger regardless of whether 2D or 3D conditions were used. However, the IS formed under 3D conditions were significantly larger than those formed on 2D gels, suggesting greater activation. These results show that T cells have larger immune synapses with APCs, become more activated, and show more proliferation when encountering antigen in a mechanically stiff 3D microenvironment.

**Figure 4.**
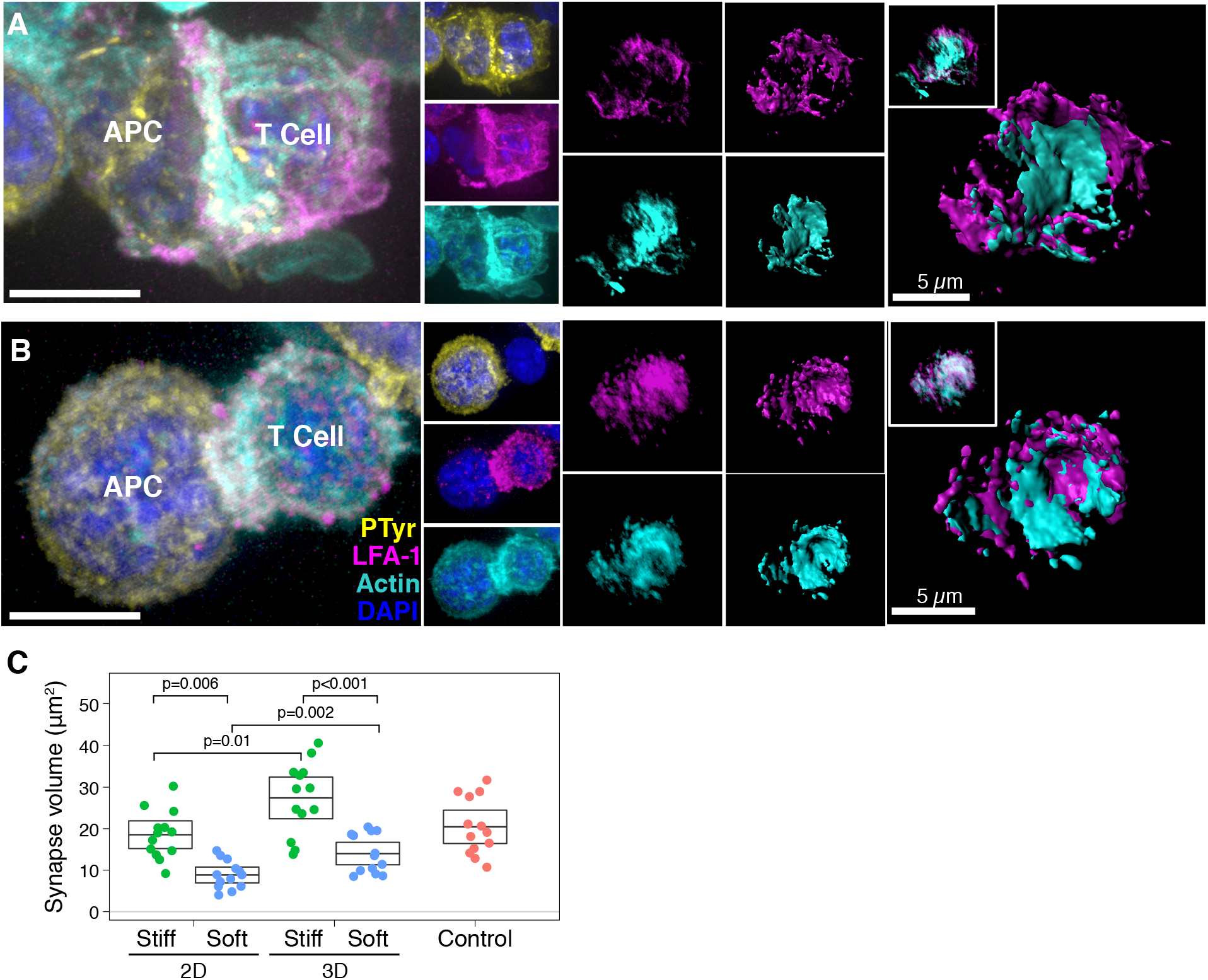
Immune synapse size is governed by the stiffness of 3D hydrogels. (**A** and **B**) Examples of IS formed by T cells and APCs seeded into **(A)** soft (4 kPa) or **(B)** stiff (40 kPa) 3D hydrogels. LFA-1 is magenta and F-actin is cyan. Cell nuclei were stained with DAPI (blue). Scale bar is 5 *μ*m. **(C)** The volume of LFA-1 at T cell-APC interfaces were measured for IS formed under 2D or 3D conditions. Control IS were formed by briefly centrifuging T cells and APCs into a pellet prior to transferring to a coverslip and fixing.

## Discussion

In vivo, T cells are activated and carry out their effector functions in a 3D tissue environment composed of polymeric matrix proteins, such as collagen, and other cells. However, the vast majority of *in vitro* experiments – which form the basis for much of our knowledge of T-cell biology – are carried out on plastic plates that are orders of magnitude stiffer than any biologically relevant substance. This practice continues even though it is accepted that the stiffness of the extracellular substrate affects the function of many different cell types (Discher et al., 2005).

Recently, several groups have demonstrated that the T-cell receptor itself senses mechanical forces through the development of a catch bond between the TCR and the pMHC (Liu et al., 2014). Our group and others have further detailed the mechanical forces acting on the TCR to trigger activation (Hu and Butte, 2016; Kim et al., 2009). We showed in fact that the forces needed to trigger the TCR arise from the stereotypical movements of the T cell itself after initial triggering. The TCR is embedded in a fluid membrane. Thus, anchoring of pMHC on a stiff substrate allows for the TCR to experience a greater force when they interact. Indeed a number of groups have investigated the role of substrate stiffness on T-cell biology using 2D synthetic polymers such as PDMS and polyacrylamide (Judokusumo et al., 2012; Saitakis et al., 2017; O’Connor et al., 2012). These groups have shown that the stiffer the anchoring, the more the T cell activation.

T cells may be able to sense mechanics through routes independent of the TCR. For example, it is well known that the intergrin LFA-1 on T cells binds to its ligand ICAM-1 on APCs. Anchoring of ICAM-1 to the APC cytoskeleton enhances the molecular forces acting on LFA-1, leading to the unfolding and “maturation” of the LFA-1 receptor into a higher affinity state for its ligand. Thus, forces acting on the T cell through the ICAM-1-LFA-1 system offer a route to costimulation that is independent of TCR triggering.

Another route by which T cells could sense forces is through microenvironmental stiffness acting on their cytoskeleton or nuclear envelope (Buxboim et al., 2017; Discher et al., 2005). This kind of mechanosensing does not act through the TCR, but rather through other cellular sensors. The role of the mechanical properties of the 3D microenvironment on T-cell activation have not been addressed. Difficulties in tuning the stiffness of 3D matrices without dramatically altering their porosity has been a roadblock in pursuing this research. The development of a freeze-fracture technique to introduce macroporosity into hydrogels of tunable stiffness represents an advance for T cell biology. Our work shows that T cells can sense their environmental mechanical environment.

## Materials and Methods

### Experimental Procedure

#### 1. Chemicals and Biologicals

Unless noted otherwise, all chemicals were purchased from Sigma-Aldrich, Inc. (St. Louis, MO). All glassware was cleaned overnight using concentrated sulfuric acid and then thoroughly rinsed with Milli-Q water. All the other cell culture reagents, solutions, and dishes were obtained from Thermo Fisher Scientific (Waltham, MA), except as indicated otherwise.

#### 2. Methods

To form the scaffolds, we first oxidized the alginate (Mw ~250 kDa, high G blocks; Novamatrix UP MVG, FMC Biopolymer, Rockland, Maine) with sodium periodate (1.5 %), overnight at room temperature, then quenched the reaction by dropwise addition of ethylene glycol (Sigma) for 45 min. We then dialyzed the solution (MWCO 3.5 kDa) against deionized water for 3 d followed by lyophilization. Afterward, the alginate was dissolved in MES (150 mM MES, 250 mM NaCl, pH 6.5) and covalently conjugated to RGD peptide (GGGGRGDY; GenScript USA Inc., Piscataway, NJ) using carbodiimide chemistry (NHS/EDC). The reaction was continued for 24 h followed by dialysis (MWCO 20 kDa) and lyophilization. This alginate-RGD complex in PBS was then cross-linked via calcium sulfate solution. The gels were casted in desired 24- or 96-well plates followed by two overnight washes to get rid of extra calcium ions and then used as 2D matrices. For 3D structures these same scaffolds were frozen at - 80 °C, lyophilized for 3 d, and stored at 4 °C before cellular studies.

We prepared an array of different alginate formulations by varying either the polymer content or the amount of crosslinker (here CaSO_4_). In order to measure the mechanical stiffness of our gels we used Instron 5542 mechanical tester and all the samples were tested at a rate of 1 mm/min. The Young’s modulus was then calculated from the slope of the linear region that corresponds with 0–10% strain. The two gels that we worked with had order-of-magnitude difference in stiffness. The softer scaffold comprised alginate 1.25% with 10 mM CaSO_4_ and had an elastic modulus of 4.1 kPa. The stiffer one comprised alginate 2.5% with 40 mM CaSO_4_ and had an elastic modulus of ~38 kPa.

Scanning electron microscopy (SEM) images of the gels were taken to see the cross-sectional microstructure and porosity of the alginate-RGD scaffolds. The lyophilized scaffolds were freeze-fractured (using liquid nitrogen) for cross-sectional images. The scaffolds were sputtered with iridium (South Bay Technology Ion Beam Sputtering) prior to imaging with a ZEISS Supra 40VP scanning electron microscope (Carl Zeiss Microscopy GmbH). The sizes of at least 40 pores from SEM images were then measured and analyzed using ImageJ software (NIH).

#### 3. Co-culture of APCs and immune cells

##### 3.1. T-cell isolation and activation

All *in vitro* experiments were conducted in strict accordance with UCLA’s institutional policy on humane and ethical treatment of animals, using five-to eight-week-old OT-II TCR transgenic mice (Jackson Labs). Cell culture media was RPMI supplemented with 10% heat inactivated FBS, 1% penicillin/streptomycin, 1% sodium pyruvate, 1% HEPES buffer, 0.1% μM 2-ME. CD4+ T cells were purified using negative enrichment kits (Stem Cell Technologies).

The common *in vitro* activation protocol for activation of CD4+ T cells was followed by culturing 1×10^6^ cells/mL in tissue culture-treated 24-well plates that were pre-coated with anti-CD3 (2C11; Bio X Cell) at a concentration of 10 μg/mL followed by addition of 2 μg/mL soluble anti-CD28 (37.51; Bio X Cell).

To form *in vitro,* 2D or 3D immune synapses we used OT-II TCR transgenic mice (Jackson Labs) were used. The I-A^b^-bearing B-cell lymphoma line LB27.4 was purchased from the American Type Culture Collection and used as APCs. T cells were extracted from spleen and CD4+ T cells were then purified using EasySep immunomagnetic negative selection (Stem Cell Technologies). Cells were then activated on plates as mentioned. Effector T cells were then collected from wells and allowed to proliferate in Interleukin-2 (IL-2, BRB Preclinical Repository, NCI, NIH) – containing medium (50 U/), prior to being used for experiments on days 3 to 5 of incubation. LB 27.4 cells were incubated with Ova peptide (1 uM) for several hours prior to co-culture with T cells. Then for control experiments in a 1.5 mL Eppendorf microfuge tube, a ratio of one B cell per one T cell were mixed, followed by centrifugation for 1.5 min at 500 x g. This pellet was then incubated at 37 °C for 15 min. Afterward the pellet was gently broken and plated in 8-well Poly-D-lysine-coated Labtek II chambers. The chamber was then centrifuged for 1 min at 50 x g. The supernatant was removed, wells were washed once with PBS and fixed with 4% PFA. Then to permeabilize cells for intracellular staining 0.1% Triton X-100 in PBS was incubated with cells for 5 min and blocked in 5% donkey serum for at least an hour. Then to stain the cells we used antibodies against LFA-1 (clone I21/7) and 4G10 with Alexa Fluor 568–conjugated phalloidin (Life Technologies) and DAPI. After several washes, wells were incubated with fluorescently conjugated secondary antibodies (Jackson ImmunoResearch). Eventually, samples were covered with Fluoromount-G with DAPI (eBioscience) and stored at 4 °C before imaging. Then synapses were measured based on LFA-1 and actin accumulation at the T cell–APC interface. For confocal imaging we used a 100× Plan Apo numerical aperture (NA) 1.4 objective (Nikon). For synapse formation on 2D or 3D hard vs. soft substrates, naive and activated CD4+ T cells were mixed with ova treated B cells the same as control experiments and then immediately seeded onto 2D or 3D matrices. Then, for experiments in a 3D context, a mild centrifugation was performed to help with dispersion of the cells within the scaffolds and incubated at 37 °C for 30 min. Afterward, to evaluate just the synapses that were formed within the scaffold and not the ones that have been formed on the plastic bottom of the wells we physically transferred the scaffolds into new wells. Following transfer, they were fixed immediately with 4% PFA for 30 min. Then the gels were lysed using EDTA (50 mM) and alginate lyase. After digestion of the hydrogel and recovery of the cells followed by three centrifugation rounds to remove the dissolved polymer. Cells then were seeded on 8 well-plate Labtek II chamber pre-modified with Poly-D-lysine. The rest of steps were followed as mentioned above.

For flow cytometry analysis, antibodies to mouse CD4, CD25 (PC61.5), CD44 (IM7), were purchased from eBioscience, BioLegend, or BD Biosciences. To study proliferation behaviour of T-cell responses during various treatments their expansion was measured by 5-(and-6)-carboxyfluorescein diacetate, succinimidyl ester (CFSE) dilution. For CFSE dilution experiments, 5 × 10^5^ naive CD4+ T cells were labeled with 2 μM CFSE for 13 min, followed by two washes and then incubation with splenocytes. Splenocytes were extracted from the spleen of wild type mice. Then the cells were incubated in ACK lysis buffer (Gibco) for 5 min at room temperature to remove red blood cells. The remaining cells were then treated with ova peptide as LB 27.4 cells prior to presentation to naive T cells. Trypan Blue were purchased from Calbiochem. Cells were analyzed on a Cytek flow cytometer using FlowJo software (Treestar).

For the experiments regarding to the velocity measurement of T-cell migration on 2D gels or within 3D scaffolds, CD4+ T cells were stained with CellTrace CFSE (1 μM) as per the manufacturer’s protocol (Life Technologies). The scaffold was mounted in a Delta T dishes (Bioptechs) which kept the media warmed to 37 °C. For experiments in presence of LB 27.4 cells as APCs, B cells were loaded with ovalbumin peptide prior to introduction to T cells and then seeded in matrices. Fields were imaged every 12 or 15 s for 10-20 min with a 40× Plan Fluor NA 0.6 extra-long working distance objective (Nikon). Imaris software (Bitplane) was used to track CellTrace CFSE-loaded T cells over time. Velocities and other statistics were exported from Imaris and analyzed in R.

#### 4. Statistical analysis

We employed permutation testing for all statistical comparisons including synapse size, proliferation, and activation markers. We used the permutationTest2 function of the “resample” package of R to calculate *P* values and determine the 95% confidence intervals, performing 50 to 100,000 permutations. Boxes in all figures show the bootstrapped mean and 95% confidence interval.

